# Differential methylation as a mediator of COVID-19 susceptibility

**DOI:** 10.1101/2020.08.14.251538

**Authors:** Sandra Steyaert, Geert Trooskens, Joris R. Delanghe, Wim Van Criekinge

## Abstract

The COVID-19 outbreak shows a huge variation in prevalence and mortality on geographical level but also within populations^1^. The *ACE2* gene, identified as the SARS-CoV2 receptor, has been shown to facilitate the viral invasion and people with higher *ACE2* expression generally are more severely affected^2, 3^. As there is a lot of variability in *ACE2* expression between individuals we hypothesized that differential DNA methylation profiles could be (one of) the confounding factors explaining this variability. Here we show that epigenetic profiling of host tissue, especially in the *ACE2* promoter region and its homologue *ACE1*, may be important risk factors for COVID-19. Our results propose that variable methylation can explain (part of) the differential susceptibility, symptom severity and death rate for COVID-19. Our findings are a promising starting point to further evaluate the potential of *ACE1/2* methylation and other candidates as a predictor for clinical outcome upon SARS-CoV2 infection.

## Background

In December 2019, a novel human coronavirus now named Severe Acute Respiratory Syndrome Coronavirus 2 (SARS-CoV2) emerged and was responsible for the now global outbreak of potentially severe and fatal atypical pneumonia defined as coronavirus disease 19 (COVID-19)^1^.

As of August 15th, 2020, SARS-CoV2 is responsible for more than 21 million detected COVID-19 cases and at least 762k deaths worldwide^4^. The outbreak of the COVID-19 pandemic is especially unnerving because it’s hard to predict how the virus will affect any individual person. As symptoms vary a lot between infected persons — if they experience them at all — determining if a person is indeed infected by this novel coronavirus is a surprisingly hard and tricky question to answer without testing^5^.

The Centers for Disease Control and Prevention (CDC) reports that the main symptoms are fever/chills, headache, muscle pain, fatigue, coughing, sore throat and shortness of breath, which can appear somewhere between two and 14 days after exposure to the virus^6^. Confirmed COVID-19 cases have shown a wide range of reported symptoms, going from mild to severe illness. According to the CDC, approximately 80% of infected people present few to mild symptoms, while others have a more severe manifestation of the disease, with extreme cases relying on a ventilator to breathe.

There are a few clear risk factors^7^, including age and general health status but even within these subgroups, there is a huge range in severity. It has been shown that next to age and health status, also socio-economic and environmental factors^8^, sex^9^, and even vitamin D status^10^ are associated with the immune response and differential susceptibility. Identifying what factors are responsible for the interindividual response to the virus is of utmost importance to aid in the identification of population groups at higher risk and to guide more effective strategies and protective measurements.

It should be noted that this difference in individual susceptibility is actually not that unusual for infectious diseases, and thus not unique for COVID-19. The same is true for tuberculosis, malaria and the ‘common’ flu^11^. The underlying reason, however, is mostly disease-specific due to the fact that the biological pathways involved in the manifestation of the illness often differ.

Certain genomic variants play a role in the specific immune response to viral infections. The last weeks, efforts were initiated to look into the role of genetics, and if each person’s unique genetics may affect their susceptibility and severity to COVID-19. Especially the interplay between the *ACE* genes and SARS-CoV2 has been of great interest.

The angiotensin-converting enzyme 2 (*ACE2*) gene, which been identified as the SARS-CoV2 receptor, has been shown to facilitate in the viral invasion^2^. While most COVID-19 studies currently focus on ACE2, a Belgian research group very recently released data from 33 European countries where they looked at the gene coding for angiotensin-converting enzyme 1 (ACE1)^12^. Although ACE2 and ACE1 share only 42% of amino acid identity, they both act as carboxypeptidases to cleave amino acids from the peptides’ carboxyl terminal. The ACE1 enzyme is characterized by a genetic deletion/insertion (D/I) polymorphism in intron 16, which is associated with alterations in circulating and tissue concentrations of ACE. Their results demonstrate that prevalence and mortality of COVID-19 infection correlate with the D allele frequency of the ACE1 D/I polymorphism. Another research group recently has launched a similar hypothesis^3^. Interestingly, the D allele has proven to be associated with a reduced expression of ACE2^3^.

It was already known that expression of *ACE2* is significantly increased amongst people who smoke or suffer from diabetes, hypertension, heart diseases and other conditions^13^, and people with higher *ACE2* expression generally seem also more severely affected by SARS-CoV2^14^. Interestingly, this mechanistically implicated *ACE2* gene has been shown to be epigenetically regulated. Taken together with the patient characteristics for severe COVID-19 outcome this substantiates an essential role for epigenetic regulation.

## Results

To explore the importance of *ACE2* methylation, publicly available DNA methylation data from 5 research studies (^~^1000 samples) was used. All data originated from blood from “healthy” people and methylation analysis was performed on Illumina Infinium HumanMethylation450 BeadChip (450K) DNA methylation arrays. As in all Illumina methylation assays, methylation values of each probe are expressed as β values, which range from 0 (no methylation) to 1 (fully methylated). Raw β values were preprocessed, cleaned and subsequently normalized. In next step, the 8 probes falling into the *ACE2* region were fetched and visualized. **Figure 1** displays for each individual 8 probe the corresponding methylation levels per sample. The x- and y-axis show each sample’s age and its normalized β value for that position/probe, respectively. Probes are sorted per genomic position in *ACE2* (left to right per row).

**Figure 1:**
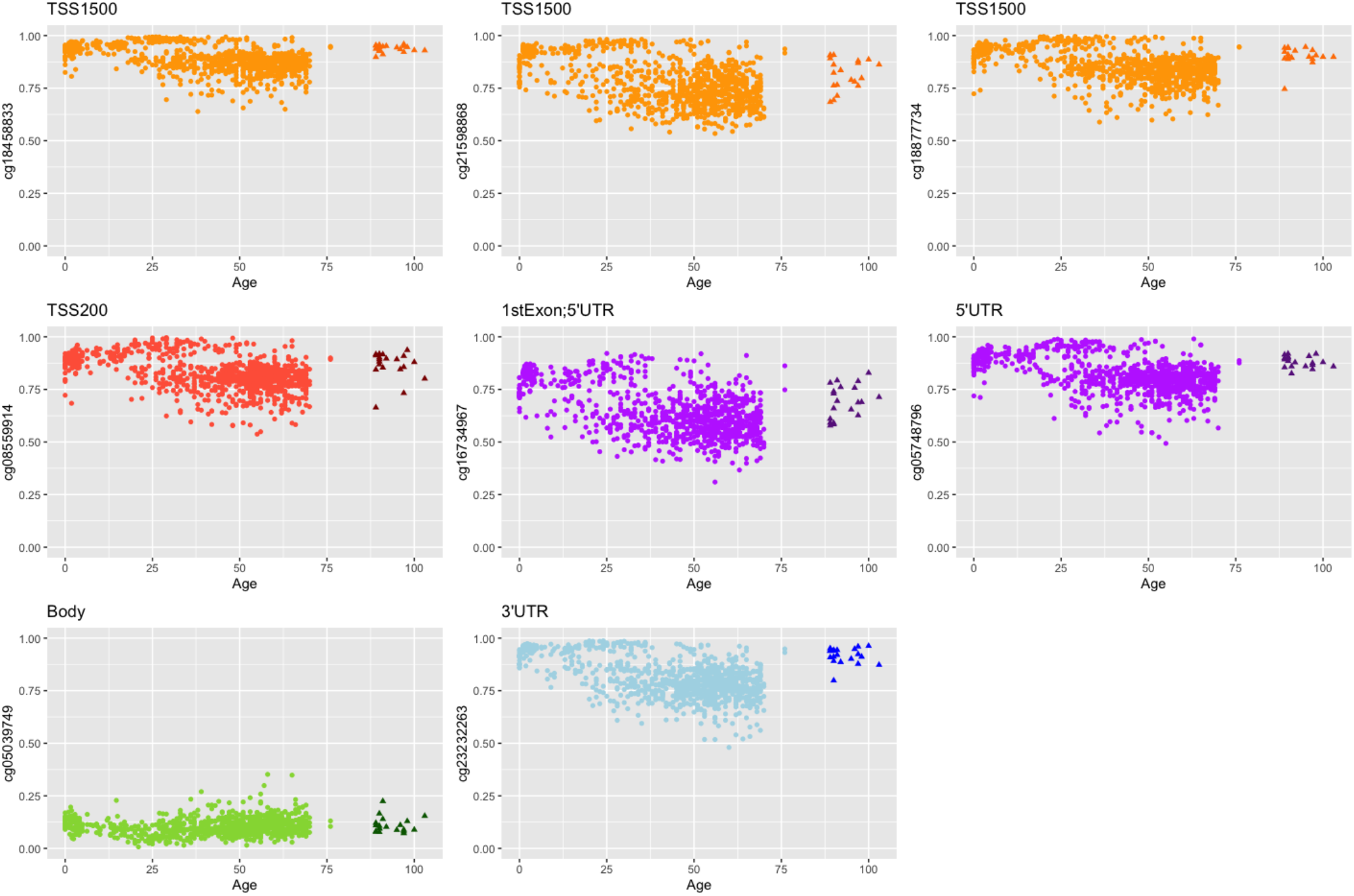
Chronological age (x-axis) vs normalized β values (y-axis) for 8 methylation probes in ACE2. Subjects older than 85years (here referred to as ‘centenarians’) are marked with a triangle and a darker color. Probes are colored as follows: orange=TSS1500; red=TSS200; purple=1stExon and/or 5’UTR; green=Gene Body; blue=3’UTR.

Two groups are noticeable: (i) the big cloud of samples marked with circles and a lighter color, and (ii) the smaller cloud of darker triangles. This latter group represents data from a cohort of centenarians. Of special interest are the promoter (flanking) regions. Here, TSS200 covers the region from transcription start site (TSS) to −200 nucleotides upstream; TSS1500 represents the region −200 to −1500 nucleotides upstream of TSS. Probes falling into these regulatory regions are colored in orange, red and purple (TSS1500/200 + 1stExon/5’UTR) It is apparent that in general upon aging the average methylation levels gradually decline while the spread increases. However, when only looking at the group of centenarians methylation levels appear higher and show a remarkable higher consistency (**Table 1**).

**Table 1:**
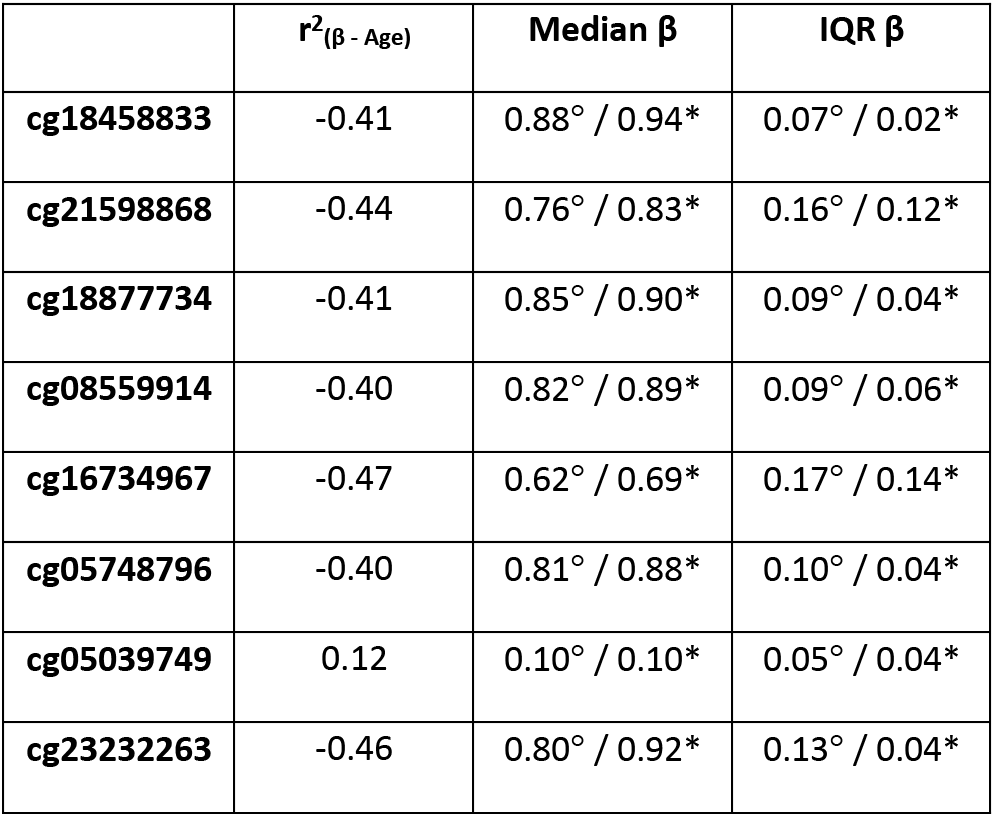
Summary statistics of methylation (β) values per probe. The first column shows the squared Spearman correlation coefficient (r^2^) between methylation and age. The second and third columns display for each probe the median β value as well as the interquartile range (IQR) for the non-centenarian group (°) and the centenarian group (*), separately.

| | $r^2(\beta - \text{Age})$ | Median $\beta$ | IQR $\beta$ |
| --- | --- | --- | --- |
| cg18458833 | -0.41 | 0.88$^\circ$ / 0.94* | 0.07$^\circ$ / 0.02* |
| cg21598868 | -0.44 | 0.76$^\circ$ / 0.83* | 0.16$^\circ$ / 0.12* |
| cg18877734 | -0.41 | 0.85$^\circ$ / 0.90* | 0.09$^\circ$ / 0.04* |
| cg08559914 | -0.40 | 0.82$^\circ$ / 0.89* | 0.09$^\circ$ / 0.06* |
| cg16734967 | -0.47 | 0.62$^\circ$ / 0.69* | 0.17$^\circ$ / 0.14* |
| cg05748796 | -0.40 | 0.81$^\circ$ / 0.88* | 0.10$^\circ$ / 0.04* |
| cg05039749 | 0.12 | 0.10$^\circ$ / 0.10* | 0.05$^\circ$ / 0.04* |
| cg23232263 | -0.46 | 0.80$^\circ$ / 0.92* | 0.13$^\circ$ / 0.04* |

There is a lot of variability in *ACE2* expression between individuals and many variants of ACE2 are associated with medical conditions such as diabetes, hypertension and cardiovascular disorders^13^. Thus, it could be that such predisposing genetic variants may also contribute to the susceptibility and explain the enormous variability in infection rate, symptom development/severity and death rate. But, while definitely plausible, to date, limited associations have been found between ACE2 variants and response to SARS-CoV2 infection. As ACE2 plays a role in the SARS-CoV2 infection pathway and *ACE2* expression seems very variable between individuals, differential DNA methylation of *ACE2* could be one of the confounding factors explaining (part of) this variability.

Hoffmann and coworkers showed that in addition to ACE2, SARS-CoV2 infection also depends on the host cell factor transmembrane protease serine 2 (TMPRSS2), a cellular protease. While ACE2 is used for initial binding and host cell entry, the spike protein of SARS-CoV2 is primed by TMPRSS2. A process of which the authors demonstrate can be blocked by a clinically proven protease inhibitor^15^. The *TMPRSS2* gene is known to be regulated by androgen^16^. Interestingly, inhibition of the androgen receptor by for example androgen deprivation therapy (ADT) decreases TMPRSS2 levels^17^ – the same effect as higher (promoter) methylation – resulting in less viral priming options and thus less viral entry^18^.

DNA methylation is not static. With age, the methylation state of various genes may change^19^. It is also known that general health status and lifestyle has an impact on the methylation signatures. These changes are quantifiable and serve as a means to determine one’s “epigenetic age” which often differs from the chronological age^20^. Epigenetic age is a collection or footprint reflecting a combination of a person’s genetic make-up but also his/her history of past external experiences/exposures. A compelling example is the case of super-centenarians (people who live > 100 years old and seem to age very healthy). When looking at their epigenetic age, it is significantly less than their chronological age. In these supercentenarians, the underlying reason is probably a combination of both genetic predisposition and lifestyle.

Could differential *ACE2* promoter methylation in host tissue be causative to the differential expression that is seen amongst COVID-19 patients? Could variable DNA signatures of *ACE2* (and *TMPRSS2*) explain (at least part of) the differential susceptibility, symptom development/severity and death rate for COVID-19? Here, we hypothesize that epigenetic profiling, especially DNA methylation signatures, may indeed permit discovery of population-, age- and gender-related risk factors for COVID-19. The more consistent methylation levels in the centenarian cohort is at least very compelling.

Our findings are a promising starting point to further evaluate the potential of *ACE2* and *TMPRSS2* methylation as a predictor for clinical outcome upon SARS-CoV2 infection. Note that blood DNA represents a mixture of DNA from various leucocyte types. However, due to close contact in the alveoli, methylation patterns found in blood can closely reflect methylation in lung tissue. Other studies focusing on the effect of smoking on DNA methylation indeed found that the effects in blood samples were very similar to the changes in lung tissue^21, 22^. But to prove any association, follow-up research is needed on respiratory samples from patients. We are now collecting these samples and aim to compare methylation profiles of *ACE2, TMPRSS2* and other interesting candidate genes from both COVID-19 negatives and positives (with ranging severity/complications). As the virus continues to spread, more mild cases will arise, and healthcare professionals need to recognize these to accurately portray total numbers of COVID-19 infections. Differentiating mild and moderate from severe disease may also help clinicians in more accurately triaging cases who need medical attention and minimize the risks on the population, health systems, and economy.

## Methods

Methylation was measured on Illumina’s 450K methylation array for which raw data was downloaded from the Gene Expression Omnibus (GEO)^23^ (GSE30870, GSE32149, GSE36064, GSE41169, GSE42861). Details on the individual data sets and characteristics of the study cohort can be found in **Table 2** below. A full description of each dataset can be found in the original reference.

**Table 2:** Details on the used 450K methylation data sets and characteristics of the study cohort.

| ID | DNA origin | Platform | #samples | Median Age (range) | Description |
| --- | --- | --- | --- | --- | --- |
| GSE30870 | Blood PMBC | 450K | 40 | 44 (0,100) | Peripheral blood mononuclear cells from newborns and centenarians. |
| GSE32149 | Blood PMBC | 450K | 46 | 15 (3.5,76) | Peripheral blood leukocytes from a DNA methylation study of Crohn's disease and ulcerative colitis. There was no significant evidence that disease status affects DNA methylation. |
| GSE36064 | Blood PMBC | 450K | 77 | 3.1 (1,16) | Leukocytes from healthy male children from Children's Hospital Boston consists of peripheral blood leukocyte samples from healthy males |
| GSE41169 | Blood WB | 450K | 94 | 29 (18,65) | Whole blood samples from a Dutch population comprised of schizophrenics and healthy control subject. It turned out that schizophrenia status was not related to DNA methylation. |
| GSE42861 | Blood WB | 450K | 689 | 54 (17,70) | Whole Blood from rheumatoid arthritis patients. Effect of rheumatoid arthritis on DNA methylation was found to be negligible. |

The Illumina 450K BeadChip measures bisulfite-conversion-based, single-CpG resolution DNA methylation levels for over 480K cytosine sites and covers 96% of CpG islands in the human genome. Unlike the previous platform (27K BeadChip) the Illumina 450K BeadChip includes two distinct probe types, Infinium I (n=135,501) and Infinium II (n=350,076). In the Infinium I type, each CpG site is targeted by two 50bp probes: one for detecting the methylated intensity (M) and one for detecting the unmethylated (U) intensity, whereas for Infinium II types, both the M and U intensity of each CpG site are detected by one single probe using different dye colors (green and red). Methylation values per CpG site are indicated by the β-value which ranges from 0 (no methylation) to 1 (fully methylated) and is computed as β=M/(M+U+α) where α is 100 generally^24^.

Raw β-values were preprocessed in R (v3.6.3) with the RnBeads package (v2.4.0)^25^. Probes not in CpG context were filtered out as well as probes for which the β values were *NA* or had low variability (standard deviation < 0.005). β-values remaining probes were next normalized using the Beta MIxture Quantile dilation method (BMIQ)^26^. In BMIQ, the β-values of type II probes are adjusted into a statistical distribution characteristic of type I probes. After normalization, probes corresponding with *ACE2* were fetched and the β-values of each sample were plotted against the subjects’ respective chronological age.

## List of Abbreviations

3’ UTR: 3’ untranslated region
5’ UTR: 5’ untranslated region
450K: HumanMethylation450 array
ADT: Androgen deprivation therapy
ACE1: Angiotensin-converting enzyme 1
ACE2: Angiotensin-converting enzyme 2
BMIQ: Beta mixture quantile
CDC: Centers for disease control and prevention
COVID-19: Coronavirus disease 19
D/I: Deletion/Insertion
GEO: Gene expression omnibus
IQR: Interquartile range
M: Methylated intensity
r^2^: squared Spearman correlation coefficient
SARS-CoV2: Severe acute respiratory syndrome coronavirus 2
TMPRSS2: Transmembrane protease serine 2
TSS: Transcription start site
U: Unmethylated intensity

## Availability of data and materials

The datasets analyzed during the current study are available in the Gene Expression Omnibus (GEO) (https://www.ncbi.nlm.nih.gov/geo/) under accession numbers GSE30870, GSE32149, GSE36064, GSE41169 and GSE42861.

## Authors’ contributions

SS pre-processed, analyzed and interpreted the methylation data and wrote the manuscript with biological and technical insight from WVC. Additional validation was done by GT. GT and JRD reviewed the analysis and final text. WVC oversaw the work and was a major contributor in writing the manuscript. All authors read and approved the final manuscript.

## Acknowledgements

We thank Dr. Adriaan Verhelle (Scripps Research, La Jolla, CA, USA) for valuable discussions, comments and help with figure layout.

